# Optimized Cas9 expression systems for highly efficient Arabidopsis genome editing facilitate isolation of complex alleles in a single generation

**DOI:** 10.1101/393439

**Authors:** Jana Ordon, Mauro Bressan, Carola Kretschmer, Luca Dall’Osto, Sylvestre Marillonnet, Roberto Bassi, Johannes Stuttmann

## Abstract

Genetic resources for the model plant Arabidopsis comprise mutant lines defective in almost any single gene in reference accession Columbia. However, gene redundancy and/or close linkage often render it extremely laborious or even impossible to isolate a desired line lacking a specific function or set of genes from segregating populations. Therefore, we here evaluated strategies and efficiencies for the inactivation of multiple genes by Cas9-based nucleases and multiplexing. In first attempts, we succeeded in isolating a mutant line carrying a 70 kb deletion, which occurred at a frequency of ~1.6% in the T_2_ generation, through PCR-based screening of numerous individuals. However, we failed to isolate a line lacking *Lhcb1* genes, which are present in five copies organized at two loci in the Arabidopsis genome. To improve efficiency of our Cas9-based nuclease system, regulatory sequences controlling Cas9 expression levels and timing were systematically compared. Indeed, use of *DD45* and *RPS5a* promoters improved efficiency of our genome editing system by approximately 25-30-fold in comparison to the previous ubiquitin promoter. Using an optimized genome editing system with *RPS5a* promoter-driven Cas9, putatively quintuple mutant lines lacking detectable amounts of Lhcb1 protein represented approximately 30% of T_1_ transformants. These results show how improved genome editing systems facilitate the isolation of complex mutant alleles, previously considered impossible to generate, at high frequency even in a single (T_1_) generation.

## Introduction

Sequence-specific nucleases (SSNs) based on SpCas9, which derives from the *Streptococcus pyogenes* CRISPR/Cas system, are currently the most commonly used tool for genome editing in plants and animals (reviewed in Ceasar et al., 2016). SpCas9 (hereafter termed Cas9) and related RNA-guided nucleases (RGNs) are directed to DNA target sequences by a guide RNA incorporating into the nuclease protein. The guide RNA may consist of a chimera from base-pairing between a CRISPR RNA (crRNA) and a trans-activating RNA (tracrRNA), or crRNA and tracrRNA may be collapsed into a single guide RNA (sgRNA; Jinek et al., 2012). In either case, the variable stretch of crRNA/sgRNA base-pairs with complementary target DNA sequences flanked by a protospacer-adjacent motif (PAM, NGG for unmodified Cas9). Cas9 functions in its native configuration as a nuclease and cleaves target sequences, but can be considered as a programmable DNA-binding scaffold for tethering diverse activities or functionalities to precise chromatin positions. As such, e.g. transcriptional regulators, chromatin modifiers or base editors were constructed on the basis of catalytically inactive Cas9 (dCas9), or fluorescent protein fusions were exploited for live cell imaging (Chavez et al., 2016; Chen et al., 2016; Dominguez et al., 2016; Komor et al., 2016; Ren et al., 2018).

In the nuclease mode, Cas9 generates either blunt-ended double-strand breaks (preferentially three base pairs upstream of the PAM sequence), or single-strand breaks when converted to a nickase (Jinek et al., 2012; Ran et al., 2013). Gene targeting may be achieved from both types of lesions, but remains technically challenging, at least in plants (Fauser et al., 2014; Shi et al., 2017). In contrast, the disruption of genes through error-prone repair of double strand breaks by non-homologous end-joining (NHEJ) is now routinely used for reverse genetics approaches in many different plant systems (reviewed in Malzahn et al., 2017). When using RGNs for precision gene editing in crop improvement, the specificity of the enzyme may be of major importance. Indeed, delivery of ribonucleoprotein complexes has been reported to minimize RGN off-target activity, and is also preferable to avoid regulation of the final product (Woo et al., 2015; Huang et al., 2016; Wolt et al., 2016; Zhang et al., 2016; Liang et al., 2017). In contrast, *Agrobacterium-facilitated* transformation and *in planta* expression yet remains the most important approach for RGN delivery in basic research. Efficiency in this case mainly depends on timing and levels of expression of nuclease and sgRNA and efficient nuclear import. Although high expression levels may increase off-target activity, these effects can be mitigated, e.g., through analysis of multiple alleles, and efficiency is generally at prime. In some species, as e.g. rice, high efficiencies for genome editing regularly approaching 100% in T_0_ plants (with Cas9 or Cpf1) were reported, suggesting that further optimization is not required in this respect (Mikami et al., 2015; Tang et al., 2017). However, especially in Arabidopsis, the genetic analyses workhorse, genome editing efficiencies are often comparably low, and severely vary between different studies. In the Arabidopsis system, transformation does not depend on somatic embryogenesis, as T-DNAs are directly delivered to female ovules during floral dip transformation (Ye et al., 1999; Desfeux et al., 2000). Accordingly, T-DNA-encoded SSNs are subsequently expressed (or not) in different cell types of the developing embryo, and expression levels and timing will be decisive for efficiency and germ line entry of genome modifications. Indeed, exceptionally high genome editing efficiencies were reported when *RPS5a* or *DD45* promoters (or derivatives) were used for Cas9 expression (Wang et al., 2015; Mao et al., 2016; Tsutsui and Higashiyama, 2017). These effects were mainly attributed to activity of these promoters in early embryogenesis. However, the cross-study comparison of genome editing efficiencies is of limited validity, as differences between constructs go beyond the use of a particular promoter. Especially the sgRNA/target site represents a major variable affecting genome editing efficiency, and also further differences between vectors may have profound and unexpected consequences.

Here, we systematically compared regulatory elements in order to determine improved Cas9 expression systems for Arabidopsis genome editing. While a previously used *ubiquitin* promoter-driven Cas9 produced mutants in the T_2_ generation and at moderate frequencies only, optimized expression systems were highly efficient in the T_1_ generation. Indeed, this enabled us to isolate a putative quintuple mutant lacking detectable amounts of Lhcb1 protein in a single generation and at high frequencies, while we had failed to isolate this mutant line prior to system optimization. This shall guide further optimization of Arabidopsis genome editing systems, and also researchers in their future choice of system.

## Material and Methods

### Plant material and transformation

Arabidopsis accession Landsberg *erecta* (L*er*), the *old3-1* mutant line (Tahir et al., 2013), accession Columbia and the NoMxB3 quadruple mutant line were used. The NoM line is published (Dall’Osto et al., 2017), and a T-DNA insertion in At5g54270 (*Lhcb3;* NASC N520342) was introgressed to generate the NoMxB3 quadruple mutant. Plants were grown in soil at a 19°C: 22°C night: day cycle (200 μE/m^2^*s, 60% relative humidity) under short (16h dark, 8h light) or long (8h dark, 16h light) day conditions. To suppress autoimmunity, plants were grown at a 26°C: 28°C night: day cycle in a growth cabinet (200μE/m^2^*s, 60% relative humidity) with either short day or long day conditions. Plants were transformed by floral dip as described (Logemann et al., 2006), and *old3-1* plants were cultivated at 28°C to suppress autoimmunity for transformation.

### Molecular cloning

Golden Gate technology (Engler et al., 2008) was used for all cloning tasks, and the Modular Cloning and Plant Parts toolkits were used (pICH/pAGM/pICSL vectors; Engler et al., 2014). Generally, 20 fmol of DNA modules were used for Golden Gate reactions with either *BsaI, BsmBI* or *BpiI* and T4 DNA Ligase. Reactions were carried out in a PCR cycler (2 min 37°C, 5 min 16°C, 10-55 cycles; terminated by 10 min 50°C and 10 min 80°C steps), and transformed either into *E. coli* TopTen or ccdB survival II cells (Invitrogen; distributed by Thermo Fisher). To generate the adaptable nuclease activity reporter (pJOG367), a linker sequence was appended to a ccdB cassette (lacking *BsmBI* and BsaI sites) by PCR amplification, and subcloned in a custom cloning vector (pJOG397) to yield pJOG395. Similarly, an amplicon of a *GUS* gene with introns (Engler et al., 2014) and a linker sequence was subcloned to yield pJOG396. These modules were subsequently assembled together with a 35S promoter (pICH51277) and an *ocs* terminator (pICH41432) in a Level 1 recipient (pICH47732) to yield pJOG367. Target sequences are inserted in this adaptable reporter scaffold as hybridized oligonucleotides by a *BsmB*I Golden Gate reaction (Online Resource 1). Promoter fragments were amplified by PCR, internal BsaI and *BpiI* restriction sites domesticated and subcloned into pICH41295. The previously reported *rbcS E9* terminator fragment (Wang et al., 2015) was amplified from *Pisum sativum* genomic DNA and subcloned into pICH41276. All genome editing constructs were assembled as previously described (Ordon et al., 2017) and essentially following the Modular Cloning strategy (Weber et al., 2011). A previously published hCas9 coding sequence (Belhaj et al., 2013) was used for assemblies, and sgRNAs were driven by a 90 nt promoter fragment of Arabidopsis U6-26, as previously described (Ordon et al., 2017). Further details and oligonucleotide sequences are available upon request. Genbank files for most important modules and nuclease vectors are provided in Online Resource 2.

### sgRNA selection and deletion screening

A local instance of CasOT (Xiao et al., 2014) was used to identify specific sgRNAs for generation of the Δ*dm2a-g* mutant. Sequence windows flanking the *DM2a* and *DM2g* genes, respectively, were defined for selection of targets, and a PacBio assembly of the Landsberg *erecta* genome was used as reference (https://www.pacb.com). Specific sgRNAs were subsequently evaluated with the sgRNA designer tool (Doench et al., 2014; Doench et al., 2016) to select those with highest predicted activity. For editing of the *DM2h* gene (promoter comparison), the target sites TGATTTCTGCTAATTCATCAAGG and TTATTGATAATAATATAGAGAGG were selected. ChopChop (Labun et al., 2016) was used for selection of target sites within *Lhcb1* genes, and potential sites were further manually curated and selected. The following target sites were used for sgRNA design: CGCGGCAGTTCGGTCCGCCAAGG [1], GCCGACCTGCCGCCTAATTGTGG [2], CACTGCAGAGATATTGAACGAGG [3], GTTATATAATGCTTGATGGATGG [4] for the Δ*dm2a-g* deletion, and GGTTCACAGATCTTCAGCGACGG [1], ATGGACCCAAGTACTTGACTCGG [2], TGTGGATAACTTCTAGCTCACGG [3], GGCTACTCAAGTTATCCTCATGG [4], GAAGCGGCCGTGTGACAATGAGG [5], AGAAGTTATCCACAGCAGGTGGG [6], GAGGACTTGCTTTACCCCGGTGG [7], AGGGGAGGAGAGAGCCATTGTGG [8] for *Lhcb1* genes. Oligonucleotides JS1382/83 (TGCAGCTGAAGATCATGGC/GACTAGCGATTGTGTCCATC) were used for detection of a Δ*dm2a-g* deletion allele. Oligonucleotides MB1/MB4 (“PCR *Lhcb1.1/.2./.3”;* A AAGCCT CTGGGTCGGTAG C A/T CTGGGTCGGTAG CC AA ACCC) and MB6/MB7 (“PCR *Lhcb1.4/.5”;* CCGGCGACTCTGTAGCCCTCA/TCCGGCGACTCTGTAGCCTTC) were used to screen for deletions at *Lhcb1* loci.

### Transgenic plant selection and estimation of genome editing efficiencies

For seed fluorescence-based selection using the FAST marker (Shimada et al., 2010), T1 or T2 seeds were spread on moist Whatman paper, and observed under a motorized SteREO Discovery.V12 microscope (Zeiss) connected to an AxioCam MRc camera. Pictures were taken under bright field conditions or UV illumination and using an RFP filter set. For selection of transgenic plants from *old3-1* transformations, seedlings were grown at 28°C, treated 3 – 4 times with BASTA and resistant plants transferred to new soil. For promoter comparison, T_1_ seedlings were transferred to 22°C, onset of autoimmunity was scored after 8d, and transgenic plants transferred back to 28°C to obtain T_2_ seeds. T_2_ seeds were directly grown at 22°C in short day conditions, and phenotypes scored 20 dag to calculate editing frequencies in individual T_2_ families. Plants were transformed with a construct conferring Hygromycin resistance (pDGE277) for generation of an *lhcb1* mutant line. Seeds were surface-sterilized, grown on MS 1/10 plates containing 0,5% sucrose, Hygromycin B (25 μg/ml) and Carbenicillin (100 μg/ml), and Hygromycin-resistant seedlings selected 15 dag.

### Gel electrophoresis, immunoblotting and sample preparation

SDS-PAGE analysis was performed with the Tris-Tricine buffer system (Schagger and von Jagow, 1987). For immunodetection, samples corresponding to 0.5 μg of Chlorophyll were loaded for each sample and electroblotted on nitrocellulose membranes. Proteins were detected with alkaline phosphatase-conjugated secondary antibodies purchased from Sigma-Aldrich (A3687). Primary antibodies used were α-Lhcb1 (AS01 004), α-PsaA (AS06 172), α-PsbB/CP47 (AS04 038) from Agrisera.

### Transient genome editing efficiency assays

For GUS-based nuclease assays, an sgRNA directed against the target site TATATAAACCCCCTCCAACCAGG was used. This target site was inserted into the adaptable reporter plasmid (pJOG367). *Agrobacterium* strains containing reporter or nuclease-encoding constructs were infiltrated alone or in a 1:1 ratio at an OD_600_ = 0.6 into leaves of four different *N. benthamiana* plants. Tissue samples of individual plants were treated as replicates, and GUS activity was determined as previously described (Ordon et al., 2017). GUS activity was normalized to the reporter alone, which was arbitrarily set to 1.

## Results

### Generation of complex alleles with *ubiquitin* promoter-driven Cas9 in Arabidopsis

We had previously developed a Golden Gate cloning-based toolkit for highly multiplexed genome editing in dicotyledonous plants (Ordon et al., 2017). In this system, expression of sgRNAs is driven by the Arabidopsis U6-26 promoter, and several different promoters were provided for Cas9 expression. Using *ubiquitin* promoter-driven Cas9 (*pPcUbi4-2*, from parsley), a 120 kb deletion encompassing the *DM2* cluster of *Resistance* genes in accession Landsberg *erecta* (Ler) was previously generated (Ordon et al., 2017). The respective Δ*dm2* deletion allele could be phenotypically selected, and occurred at low frequencies only (~0.5%) among T2 individuals. To evaluate strategies for and feasibility of generating large deletion alleles not linked to a phenotype, we attempted deletion of ~ 70 kb within the *DM2*^L*er*^ cluster containing all genes of the cluster except *DM2h* (Fig. 1a). The *DM2*^L*er*^ cluster encodes 7-8 complete or truncated *Resistance* genes (*DM2a-h*, or *RPP1-like^Ler^ R1-R8*) most similar to *RPP1*, conferring resistance to the obligate biotrophic oomycete *Hyaloperonospora arabidopsidis* in accession Wassilijewska (Botella et al., 1998; Alcazar et al., 2009; Chae et al., 2014). The function(s) of the *DM2*^L*er*^ locus remain yet unknown, but one or several *DM2*^L*er*^ genes provoke constitutive activation of immune responses (autoimmunity) when combined in a single genetic background with different “inducers”: Alleles of *STRUBBELIG-RECEPTOR FAMILY 3* originating from the South Asian accessions Kashmir and Kondara (SRF3^Kas/Kond^; Alcazar et al., 2010), a transgene encoding for ENHANCED DISEASE SUSCEPTIBILITY1 fused to YFP and an SV40 nuclear localization signal (EDS1-YFP^NLS^; Stuttmann et al., 2016) or the *onset of leaf death3-1* allele affecting a cytosolic O-acetylserine(thiol)lyase (*old3-1;* Tahir et al., 2013). *The DM2h* gene is required for all cases of autoimmunity, but the contribution of additional *DM2* genes remains unknown (Alcazar et al., 2014; Stuttmann et al., 2016). The Δ*dm2a-g* deletion was conducted in the L*er old3-1* background, and plants were grown at 28°C to suppress temperature-sensitive autoimmunity and seedling necrosis in this line (Tahir et al., 2013).

**Fig. 1:**
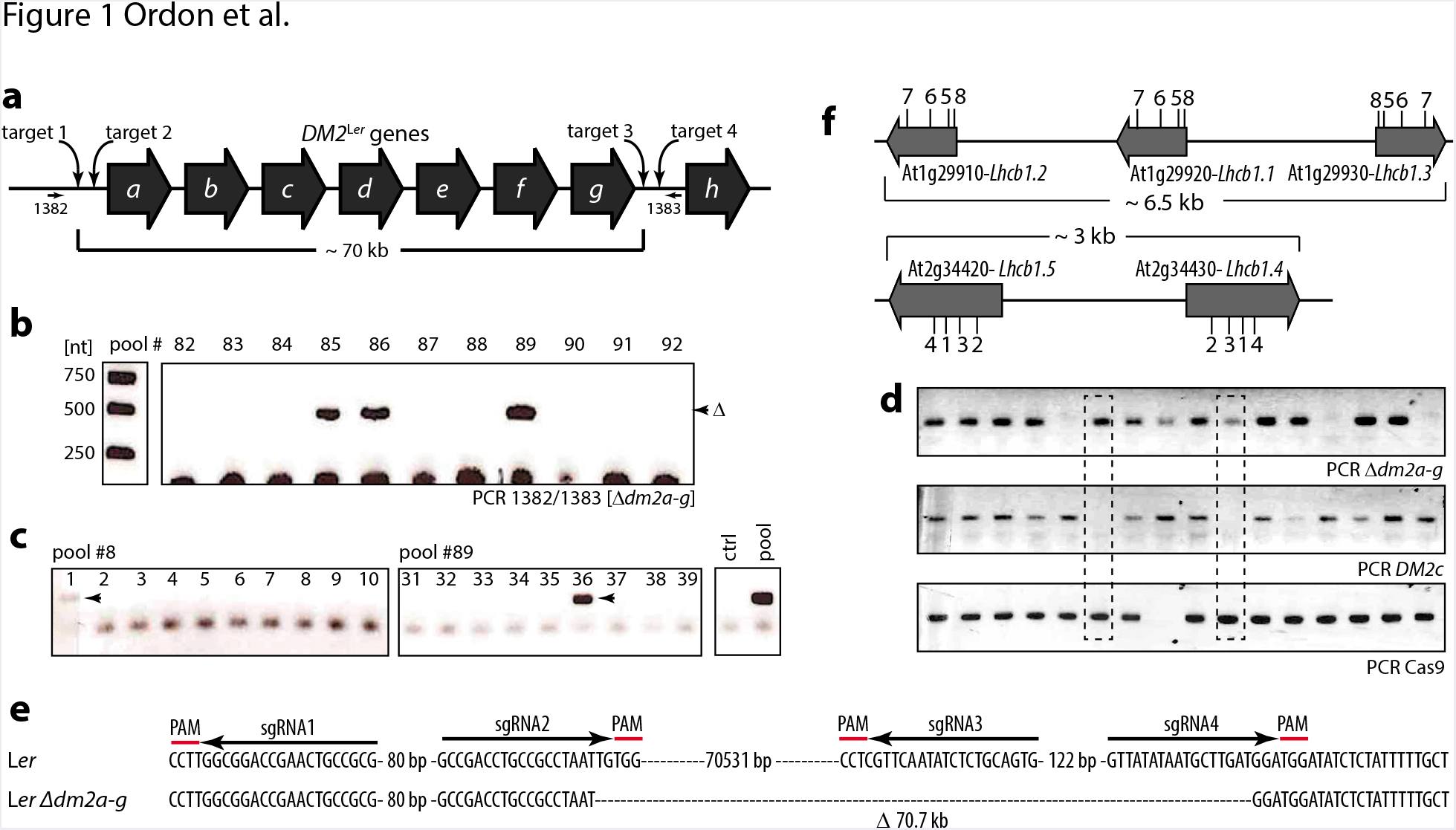
*Ubiquitin* promoter-driven Cas9 for generation of complex alleles. (a) Schematic drawing of the *DM2* cluster from accession Landsberg *erecta* (not drawn to scale). The location of sgRNA target sites and PCR primers for screening (1382/1383) is indicated. (b) PCR-screening of pooled DNAs for occurrence of a Δ*dm2a-g* allele. Each pool contained 7-11 T_2_ individuals from transformation of the Δ*dm2a-g* genome editing construct. A size of ~ 500 bp is expected upon deletion of *DM2a-g.* PCR products were resolved on a 1% agarose gel and DNA visualized with ethidium bromide. (c) Deconvolution of pools to identify single plants carrying the Δ*dm2a-g* deletion allele. DNA was extracted from single plants of pools #8 and #89 from (b), and PCR-screened for a Δ*dm2a-g* deletion as before. The parental line (ctrl) and a previously PCR-positive pool DNA were used as controls. (d) Inheritability and segregation of the Δ*dm2a-g* deletion allele in the T_3_ generation. DNA was extracted from single T_3_ plants derived from plant #36 in (c), and was used for genotyping: Presence of Δ*dm2a-g*, presence of *DM2c*, and presence of Cas9. Results shown are representative for several independent T_3_ populations analyzed in parallel. Two individuals homozygous for Δ*dm2a-g* (absence of *DM2c*) are boxed. (e) Molecular lesion in a Δ*dm2a-g* deletion line. The amplicon from (d) representing the Δ*dm2a-g* deletion was sequenced directly. The sgRNA target sites are indicated. (f) Genetic organization of the two *Lhcb1* linkage groups on chromosome 1 and chromosome 2 of the Arabidopsis genome (drawn to scale). 1-8 indicate the positioning of sgRNA target sites.

For deletion of *DM2a-g*, a construct containing Cas9 driven by the pPcUbi4-2 promoter and four sgRNAs directed against sites flanking the targeted region was transformed into L*er old3-1* plants (Fig. 1a). T_1_ plants were selected by BASTA resistance, and T_2_ populations consisting of 4-5 T_1_ plants assembled. 96 DNA pools containing 7-11 T_2_ plants each (equivalent of ~ 850 T_2_ plants) were screened by PCR with oligonucleotides flanking the targeted region for occurrence of a Δ*dm2a-g* deletion allele (Figs. 1a,b). A clear signal of the expected size was detected in 14/96 DNA pools, representing 15% of pools or approximately 1.6% of T_2_ plants considering a single line with a deletion allele in each PCR-positive pool. Selected pools were deconvoluted, and single plants screened (Fig. 1c). For some pools, only a weak signal corresponding to the deletion allele was detected among single plants (pool #8 in Fig. 1c), but most pools contained 1-2 plants positive for the deletion allele (pool #89 in Fig. 1c). T_3_ seeds were obtained for these single plants, and PCR-screened for segregation of the Δ*dm2a-g* allele and Cas9. From analysis of 96 T_3_ plants, several lines homozygous for the Δ*dm2a-g* allele (absence of *DM2c*, boxed lanes in Fig. 1d) could be isolated, but all still contained the genome editing transgene, as detected by presence of Cas9. Outcrossing of the Cas9 construct is most likely not required for most experimental settings. However, sequencing of the Δ*dm2a-g* deletion allele in one of the isolated homozygous lines revealed that it still contained an intact sgRNA target site (Fig. 1e), suggesting it might not be fully stable in subsequent generations. Summarizing, PCR-based isolation of large deletion alleles is feasible with the used genome editing system (containing pPcUbi-driven Cas9), but deletions occur at low frequency, and isolation requires extensive screening.

Furthermore, we attempted to generate a mutant lacking Lhcb1 (chlorophyll a/b binding protein 1, CAB), a major subunit of light-harvesting complex II (LHCII) and the most abundant membrane protein in nature (Galka et al., 2012; Su et al., 2017). The *Arabidopsis thaliana* genome contains five *Lhcb1* genes, which are organized in tight linkage groups on chromosomes 1 and 2 (Fig. 1f). For inactivation of *Lhcb1* genes, a genome editing construct (with pPcUbi4-2-driven Cas9) containing eight sgRNAs was constructed, and transformed into Columbia (Col) wild type plants. *Lhcb1* genes in each linkage group share high sequence homology, and sgRNAs were designed to target four sites with perfect match in each *Lhcb1* gene (Fig. 1f). Thus, genome editing activity might produce SNPs within individual genes, or larger deletions between target sites. As for Δ*dm2a-g*, T_2_ plants were PCR-screened for occurrence of larger deletions with flanking oligonucleotides. From screening ~ 200 T_2_ plants, no line PCR-positive for a deletion in either of the linkage groups could be detected. Also, none of ~ 400 plants visually inspected showed the pale green phenotype expected for lines with reduced Lhcb1 levels (Pietrzykowska et al., 2014). We concluded that efficiency of our genome editing system was not sufficient for convenient isolation of complex or highly multiplexed alleles.

### Improved genome editing efficiencies through optimized Cas9 expression systems

To improve efficiency of our genome editing system, we focused on regulatory elements controlling Cas9 expression. The *rbcS E9* terminator from pea (*Pisum sativum*) was previously described to positively affect genome editing efficiencies in comparison to the *nos* terminator (*nopaline synthase; Agrobacterium tumefaciens*) in several independent constructs and transformations (Wang et al., 2015), suggesting stabilization of the respective mRNA. To more generally test the importance of transcriptional terminators for Cas9 activity, we wanted to compare genome editing efficiencies of nuclease constructs differing only in 3’UTR sequences and transcriptional terminators for Cas9 expression. Since mRNA stabilization should not be strictly limited to a particular plant system, genome editing efficiencies were compared in quantitative, transient assays in *Nicotiana benthamiana* (*N. benth*.). For this, an adaptable genome editing efficiency reporter was first constructed (Online Resource 1). This adaptable reporter allows insertion of user-defined target sequences in a linker region of a *β-glucuronidase* (*uidA*) gene. Insertion of a target sequence disrupts the *GUS* reading frame, which can be restored by RGN-mediated cleavage and repair through the NHEJ pathway (Online Resource 1). Transient co-expression of reporter constructs and corresponding nucleases in *N. benth.* faithfully restored GUS activity, and reporter/nuclease combinations showed variable GUS activities potentially reflecting sgRNA efficacy (Online Resource 1). Nine minimalistic nuclease constructs containing only the sgRNA transcriptional unit and 35S promoter-driven Cas9 with different transcriptional terminators were quantitatively compared for genome editing efficiency (Fig. 2a). Across multiple biological replicates, no significant and reproducible differences were measured (Fig. 2b). However, high genome editing efficiencies were obtained in all replicates when using the *rbcS E9* terminator. This corroborates the previous notion that the *rbcS E9* terminator is well-suited for Cas9 expression even when combined with hCas9 (human codon-optimized), in contrast to previously used zCas9 (*Zea mays* codon-optimized; Wang et al., 2015). Furthermore, this is in line with potential masking of the beneficial effects of this terminator by expression of Cas9 by strong constitutive promoters (Wang et al., 2015). Since no other terminator out-competed *rbcS E9*, this was used for further experiments.

**Fig. 2:**
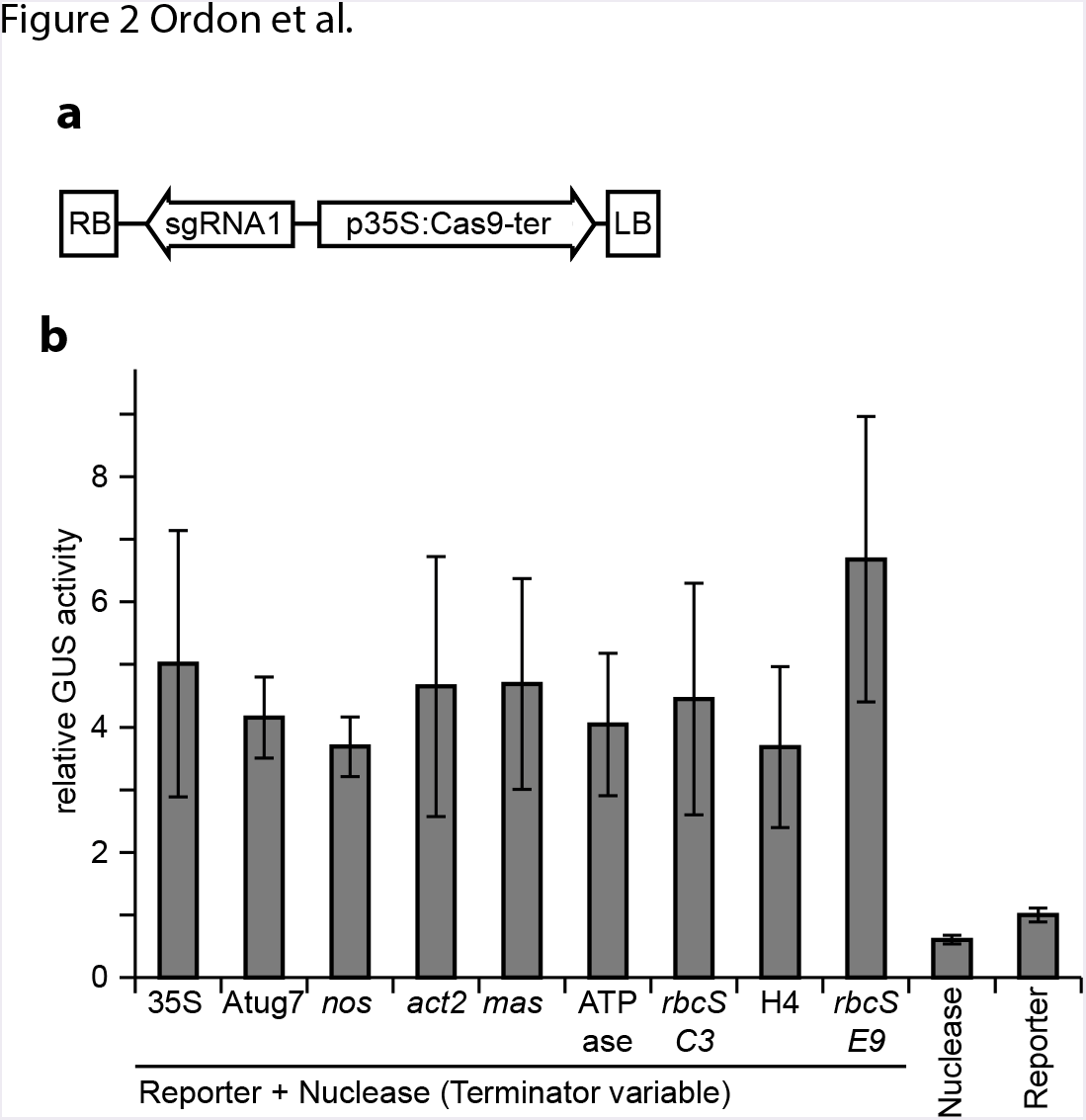
Effect of transcription terminators on Cas9 activity in transient reporter assays. (a) Architecture of minimalistic nuclease constructs used for systematic comparison of 3’UTR sequences and transcriptional terminators for Cas9 expression. (b) Evaluation of genome editing efficiency of nuclease constructs differing only in Cas9 3’UTR/terminator sequences. A GUS-based nuclease activity reporter and different nuclease constructs were transiently cotransformed into *N. benth.* tissues by Agroinfiltration, and GUS activity was determined 3 dpi. Background activity of the reporter alone was arbitrarily set to 1, and GUS activities normalized. A nuclease construct with t35S-terminated Cas9 was included as control (nuclease). Error bars represent standard deviations from four replicates. The experiment was repeated four times with similar results. The following 3’UTR/terminator sequences were used: 35S - 35S terminator from cauliflower mosaic virus; Atug7 - Atug7 terminator from *Agrobacterium tumefaciens (A. tumefaciens); nos* - *nos* terminator from *A. tumefaciens; act2 - Actin2* terminator from *Arabidopsis thaliana; mas - mas* terminator from *A. tumefaciens; ATPase - ATPase* terminator from *Solanum lycopersicum (S. lycopersicum); rbcS C3* - from *S. lycopersicum;* H4 - *Histone H4* from *Solanum tuberosum* (all Engler et al., 2014, and references therein); *rbcS E9* - from *Pisum sativum* (Wang et al., 2015).

Next, different promoters were tested for their effect on genome editing efficiency. As far as we are aware, the 35S promoter (e.g. Feng et al., 2013; Ordon et al., 2017), several different *ubiquitin* promoters (e.g. Mao et al., 2013; Fauser et al., 2014; Peterson et al., 2016), egg cell-specific promoters or derivatives (*DD45* and EC1.2-EC1.1; Wang et al., 2015; Mao et al., 2016), promoters of *APETALA1* (*AP1;* Gao et al., 2015), *INCURVATA2* (*ICU2;* Hyun et al., 2015), *YAO* (Yan et al., 2015), *SPOROCYTELESS* and *LAT52* (Mao et al., 2016), MGE1/2/3 (Eid et al., 2016), *HISTONE H4* and *EF1α* (Osakabe et al., 2016) and *RIBOSOMAL PROTEIN S5a* and *WUSCHEL RELATED HOMEOBOX 2* (*RPS5a* and *WOX2;* Tsutsui and Higashiyama, 2017) were previously used to drive Cas9 expression for Arabidopsis genome editing. The occurrence of homozygous mutants in the T_1_ generation at high frequencies was mainly reported for the *RPS5a* promoter and the egg cell-specific *DD45* promoter or derivatives (Wang et al., 2015; Mao et al., 2016; Tsutsui and Higashiyama, 2017), and also for the meiosis-specific MGE1 promoter (Eid et al., 2016). We decided to focus on the *RPS5a* and *DD45* regulatory elements, and to compare these with several popular promoters not reported to generate mutations in T_1_ generation (p35S, pPcUbi, *pAP1, pICU2*) and two further promoters not previously employed for genome editing (*pGILT, pALB;* see Online Resource 2 for sequence details for used promoter fragments).

Promoter fragments were used for driving Cas9 expression (terminated by *trbcS E9*) in genome editing vectors containing two sgRNA transcriptional units, a BASTA resistance cassette and also the FAST marker for seed coat fluorescence-based identification of transgenic plants (Fig. 3a; Shimada et al., 2010). The positioning and orientation of the FAST cassette was altered in comparison to vectors we had previously generated and containing this element (Ordon et al., 2017). Although transgenic plants could be selected by monitoring seed fluorescence with the previous vector architecture, the antibiotic/herbicide resistance markers neighboring the FAST cassette were not functional, and no genome editing activity was observed in several independent experiments and with different sgRNAs in Arabidopsis. This shall act as a cautionary note for use of the FAST marker in novel assemblies, and exemplifies the synthetic biology crux that a system’s performance is not simply the sum of its components. With the novel vector architecture, transgenic plants could conveniently be selected by herbicide resistance in the T1 generation (Fig. 3b), and also the FAST marker was functional for selection/counter-selection in T1 and T2 generations (Fig. 3c). The eight constructs differing only in promoter fragments driving Cas9 expression contained sgRNAs for targeting of the *DM2h* gene of the *DM2*^L*er*^ cluster, and were transformed into L*er old3-1* plants (cultivated at 28°C to suppress autoimmunity). Inactivation of *DM2h* rescues the autoimmune phenotype of the *old3-1* line in a dose-dependent manner, and both heterozygous (*DM2h/dm2h*) and homozygous (*dm2h/dm2h*) plants can be phenotypically identified by simple survival at different temperature regimes (Ordon et al., 2017).

**Fig. 3:**
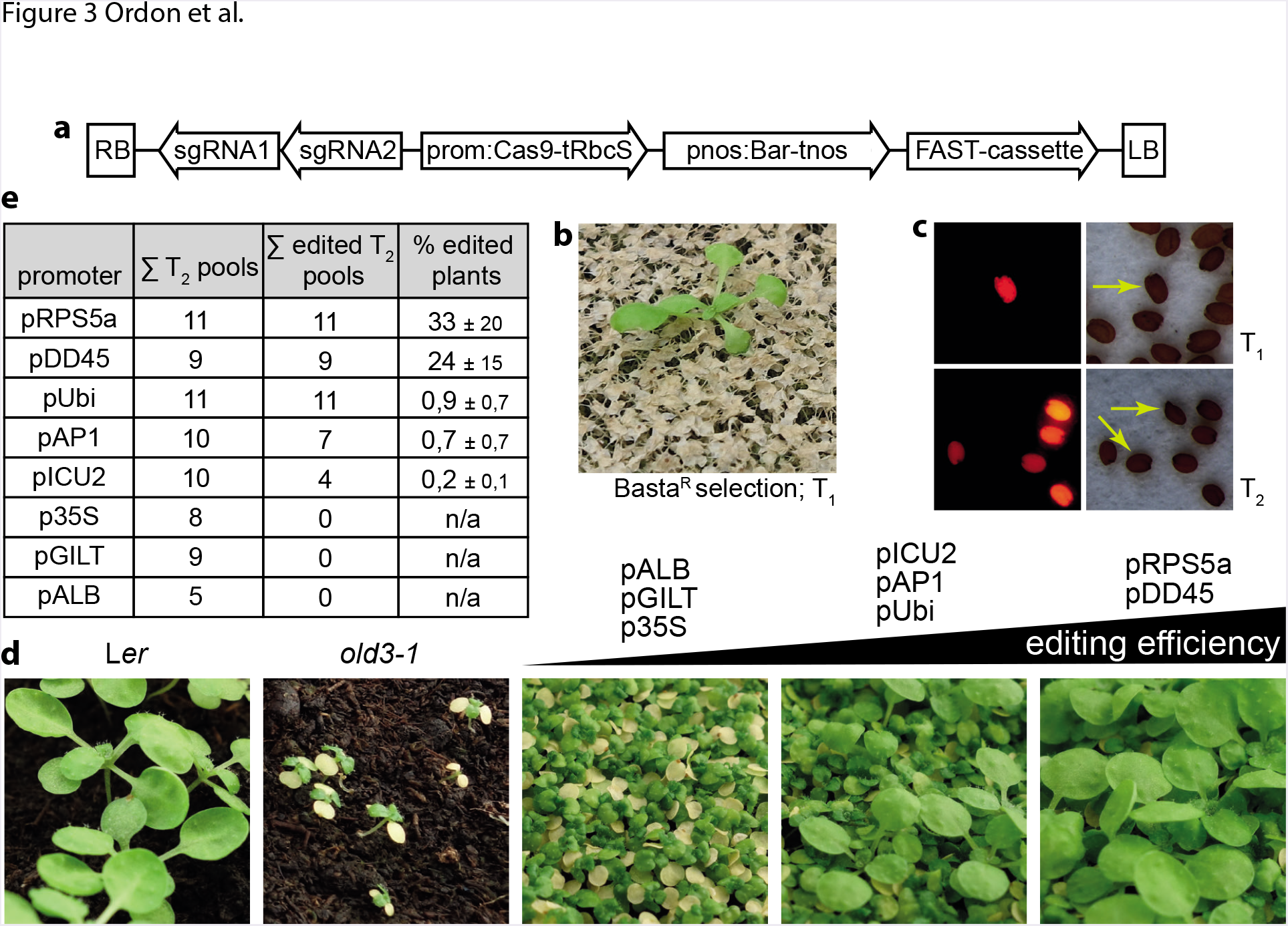
Systematic comparison of promoters for driving Cas9 expression in *Arabidopsis thaliana*. (a) Schematic drawing of constructs used for Arabidopsis transformation. Both sgRNAs are driven by identical fragments of the pU6-26 promoter. Constructs differ only in Cas9 promoter/5’UTR sequences. (b) Functionality of the BASTA resistance marker. (c) Functionality of the FAST marker in T_1_ and T_2_ generations. (d) Phenotypic survey of genome editing efficiencies with different promoters driving Cas9 expression. Representative pictures of T_2_ pools and control plants (Ler, L*er old3-1*) grown at 22°C are shown. (e) Quantitative assessment of genome editing efficiencies. Necrotic/rescued plants from (d) were counted.

Genome editing in the T_1_ generation was first tested by shifting BASTA-resistant transformants from 28°C to 22°C. At this temperature, inactivation of a single *DM2h* allele is sufficient to suppress autoimmunity of the parental line. Eight days after shifting, all plants showed signs of autoimmunity, arguing against efficient genome editing with any of the constructs/promoters in this generation and with the used sgRNAs. Five to eleven BASTA-resistant T_1_ plants of each transformation were further cultivated at 28°C to obtain T_2_ seeds. T_2_ plants were grown alongside control plants (Ler, L*er old3-1*) at 22°C, and *old3-1* plants became necrotic after 20 d of growth (Fig. 3d). Although only limited numbers of T_2_ pools were analyzed for constructs containing different promoters, obvious differences for the frequencies of phenotypically rescued (Ler-like), and thus genome-edited, plants became evident: Rescued plants were not present upon expression of Cas9 by 35S, *ALB* or *GILT* promoters, were rare for *ICU2, AP1* and *Ubi* promoters, and frequent with *pDD45* or *pRPS5a* driving Cas9 expression (Fig. 3d). We further quantified efficiencies by counting rescued plants across T_2_ populations (Fig. 3e). When comparing *Ubi, AP1* and *ICU2* promoters, similar genome editing efficiencies of approximately 1% were observed among T_2_ plants. Nonetheless, the *Ubi* promoter might perform somewhat better, since all analyzed T_2_ populations contained rescued, non-necrotic plants. In clear contrast, rescued plants occurred in all T_2_ batches from transformation of *pRPS5a-* and pDD45-containing constructs at high frequencies averaging to 33 and 24%, respectively (Fig. 3e). Up to 70% of rescued plants were observed in some T_2_ populations, but all still contained necrotic plants, confirming that no homozygous mutants were obtained in the T_1_ generation in our experiments. The differences between *RPS5a* and *DD45* promoters observed in our comparison were not statistically significant. Summarizing, our promoter comparison clearly points out superior performance of the *RPS5a* and *DD45* promoters for Arabidopsis genome editing. It should be noted that performance of *pDD45* should be further enhanced by use of the derived “EC1.2-EC1.1 fusion promoter”, which was not tested here (Wang et al., 2015).

### High frequency generation of quintuple mutants in T_1_ generation with improved Cas9 expression system

Having identified regulatory elements suitable for high efficiency Arabidopsis genome editing, the generation of an *lhcb1* mutant line was reattempted. A genome editing construct similar to that shown in Fig. 3a, but containing pRPS5a-driven Cas9, a Hygromycin resistance marker cassette (*pnos:hpt-tnos;* instead of pnos:Bar-tnos) and the same eight sgRNAs previously used (Fig. 1f) was transformed into a NoMxB3 mutant line already defective in *Lhcb4.1, Lhcb4.2, Lhcb5* and *Lhcb3* genes. T_1_ plants were selected by Hygromycin resistance, and further grown in soil. From 30 Hygromycin-resistant plants, 20 showed clearly reduced chlorophyll accumulation in comparison to the parental line, and thus the pale green phenotype expected for reduced *Lhcb1* function (Pietrzykowska et al., 2014). From the 20 plants phenotypically distinct to the NoMxB3 line, 10 had an intermediate phenotype, and the remaining 10 were severely pale (Fig. 4a). To evaluate remaining levels of Lhcb1, leaf tissue samples of three severely pale T_1_ plants and the parental NoMxB3 line were used for immunoblot analysis (Fig. 4b). While the control proteins PsaA (PSI core subunit; Mazor et al., 2017) and PsbB/CP47 (PSII core subunit; Wei et al., 2016) were detected to similar levels in all lines, no signal was obtained for Lhcb1 in the genome edited T_1_ individuals, suggesting that all five *Lhcb1* genes had been inactivated. The accumulation of light harvesting complex II (LHCII), of which Lhcb1 is a major subunit, was further analyzed by SDS-PAGE and Coomassie staining. Intensity of the major band corresponding to LHCII was strongly reduced in comparison to the parental NoMxB3 line, and we assume that the residual LHCII signal in *lhcb1* genome edited lines originates exclusively from Lhcb2, another LHCII subunit (Liu et al., 2004). PCR-screening with oligonucleotides flanking regions targeted for deletion or mutagenesis indicated complex and diverse rearrangements at the *Lhcb1* loci in putative mutant lines (Fig. 4d). In all lines, novel deletion alleles not present in the parental line (ctrl, NoMxB3) were detected. The amplification of multiple PCR products (> 2) from individual lines might result from detection of multiple somatic events (chimeric mutants), or also from low specificity of oligonucleotides due to high homology within *Lhcb1* genes. Analysis of transgene-free T_2_ individuals will be required to reveal the precise molecular lesions in putative *lhcb1* lines, and this analysis is yet ongoing. Nonetheless, these results suggest that the improved Cas9 expression system not only enhances overall genome editing frequencies, but even allows the isolation of complex alleles, in this case a quintuple mutant, in a single step in the T_1_ generation.

**Fig. 4:**
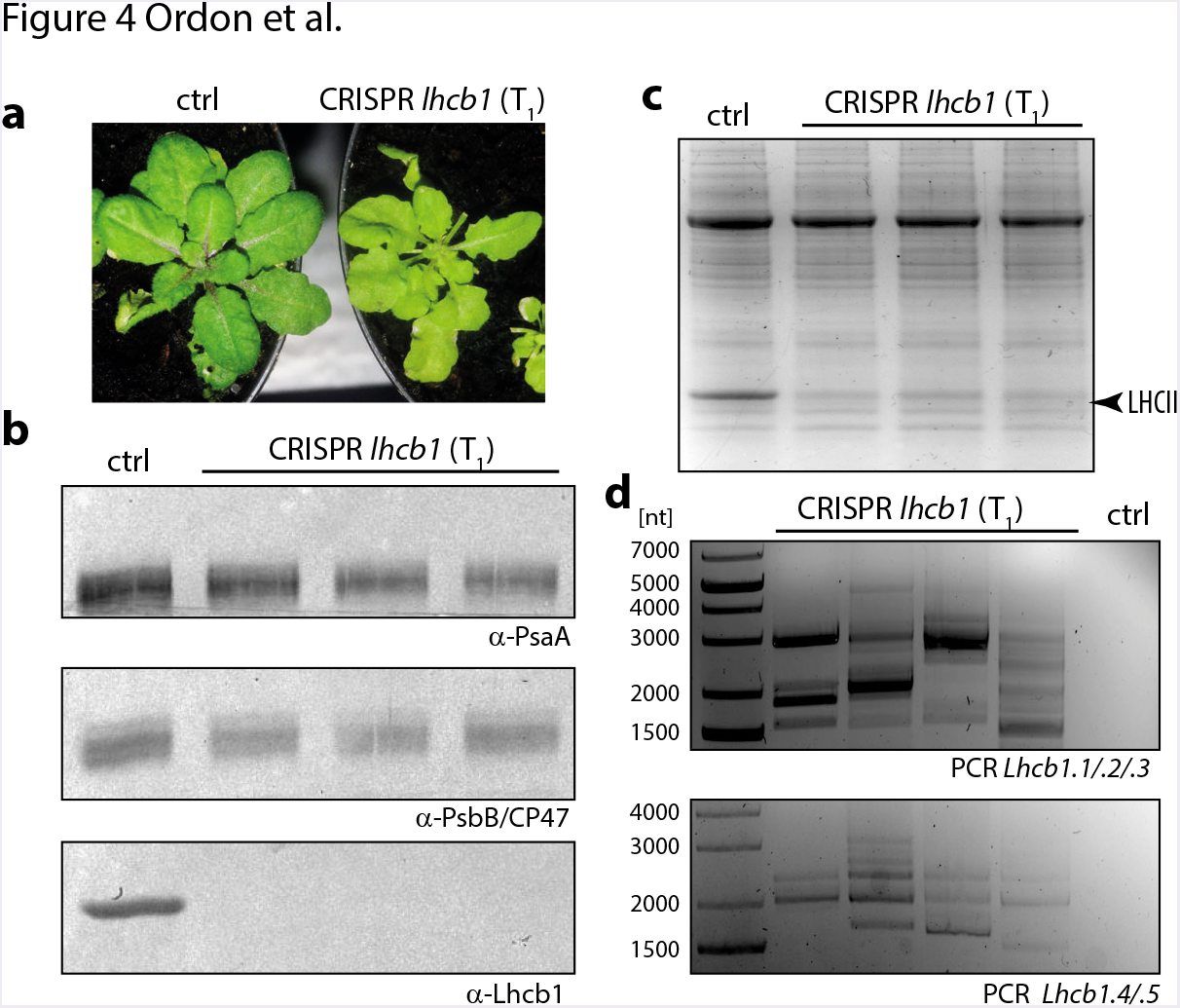
Generation of *Lhcb1* mutant plants in a single generation. (a) Phenotype of putative *lhcb1* mutant plants. The NoMxB3 parental line is shown as control, in comparison to one of the severely pale T_1_ lines recovered from Hygromycin selection and editing of *Lhcb1* genes. (b) Immunoblot analysis of Lhcb1 protein accumulation. Protein extracts from the parental NoMxB3 line and three independent T_1_ plants putatively deficient in *Lhcb1* genes were used for immunodetection of Lhcb1. PsaA and PsbB/CP47 were detected as control proteins and loading control. (c) Tris-Tricine SDS-PAGE and Coomassie staining of total protein as in (b). The major signal corresponding to LHCII is marked, and strongly reduced in genome-edited T_1_ individuals due to loss of Lhcb1. (d) PCR interrogation at *Lhcb1* loci in T_1_ genome-edited lines. DNA was extracted from T_1_ lines and the parental NoMxB3 line (ctrl), and used for PCR with primers flanking outermost sgRNA target sites (Fig. 1f) in *Lhcb1* genes on chromosome 1 (upper panel, PCR *Lhcb1.1/.2./.3*) and chromosome 2 (lower panel, PCR *Lhcb1.4/.5*).

### Discussion

Until recently, higher order Arabidopsis mutants could be generated exclusively by crossing of lines harboring the respective lesions. A number of e.g. quintuple, hexuple or even up to decuple mutants were previously reported (e.g. Fujii et al., 2011; Maekawa et al., 2012; Wild et al., 2016), but their isolation is extremely laborious due to segregation of multiple alleles and/or close linkage between loci of interest. Accordingly, it was previously stated that due to “close genetic linkage, loss-of-function (quintuple *Lhcb1* or triple *Lhcb2*) T-DNA KO mutants are almost impossible to generate” (Pietrzykowska et al., 2014). The application of SSNs as reverse genetics tools theoretically alleviated these limitations, but commonly suffered from low efficiency in first reports. Here, by using an optimized Cas9-based genome editing system with high multiplexing capacity, we show that complex alleles such as higher order (quintuple) mutants may be generated with high efficiency even in a single generation (T_1_). This demonstrates how SSNs can, through current and future optimization steps, match the increasingly complex demands and requirements of basic research projects in the Arabidopsis model system. The finding that the egg-cell specific promoters (*DD45* and especially the EC1.2-EC1.1 fusion promoter) or the *RPS5a* promoter confer particularly high genome editing efficiencies readily in the T_1_ generation (Wang et al., 2015; Tsutsui and Higashiyama, 2017) is by itself not new, but is shown here to withstand direct comparison with other promoter systems using identical vector architectures and sgRNAs/target sites. Vector maps and sequence details for optimized vectors also providing positive/negative selection used here (Fig. 3) are provided in Online Resource 2. Vectors allow simple Golden Gate-based assembly of multiplexing constructs containing up to eight sgRNAs in four days without any PCR steps as previously described (Ordon et al., 2017), and are available upon request.

The expression of Cas9 under control of the *DD45* or *RPS5a* promoters improved genome editing efficiencies at the *DM2h* locus approximately 25-fold in comparison to *ubiquitin, AP1* and *ICU2* promoters (Fig. 3). Nonetheless, we were successful in isolating a 70 kb deletion allele produced by the previous, non-optimized system containing *ubiquitin* promoter-controlled Cas9 (Fig. 1). This validates the used strategy of PCR-screening large numbers of pooled T_2_ individuals and may act as guidance for future isolation of deletion alleles for functional interrogation of gene clusters or non-coding regions. However, this strategy is obviously hampered by lacking controls for functionality of the conducted PCR prior to detection of the desired deletion allele. Interestingly, we estimated the occurrence of the 70 kb deletion allele to approximately 1.6% of T_2_ individuals, but detected editing at the *DM2h* locus (in promoter comparison experiments, Fig. 3) among only 1% of T_2_ individuals under similar Cas9 expression conditions. Our previous data suggested that point mutations were the most frequent type of Cas9-induced alleles in Arabidopsis, and that the frequency of deletion alleles (between sgRNA target sites in multiplexing applications) was inversely correlated with deletion size, as also reported in at least some studies from animal systems (Canver et al., 2014; Ordon et al., 2017). Taken together, this suggests poor efficiency of the sgRNAs used for *DM2h* editing, which may also explain failure to isolate hetero- or homozygous *dm2h* mutants in the T_1_ generation when using *DD45* or *RPS5a* promoters. Indeed, mutations in the T_1_ generation were also rare when the *GLABRA2* locus was targeted with DD45-driven Cas9 (Mao et al., 2016), supporting that recovery of Ti-edited plants might strongly depend on sgRNAs and/or target sites. Notably, all of the Δ*dm2a-g* deletion alleles detected by PCR screening and further analyzed were heritable, and segregated at Mendelian ratios in respective T_3_ generations (Fig. 1d), suggesting that detection of somatic genome editing events is not a major issue for isolation of deletions at least under the used conditions.

An optimal genome editing system for reverse genetics in Arabidopsis will produce homozygous mutants at near 100% efficiency in the T_1_ generation at any given locus. Although the optimization of Cas9 expression conditions tremendously improved efficiencies, further modifications are required to obtain this goal. In respect to Cas9 regulatory sequences, we here focused on previously reported elements, and it is well conceivable that yet uncharacterized promoters and/or transcriptional terminators might further improve genome editing efficiencies. However, also all remaining components may be further optimized, and additional functionalities implemented into T-DNA constructs. To this end, e.g. nuclear import of Cas9 might be enhanced by different or additional nuclear localization signals, or its expression improved by codon optimization or addition of introns. Furthermore, especially sgRNA expression levels appear to have a major influence on a system’s performance. To date, sgRNAs were mainly expressed directly from Polymerase III (Pol III)-transcribed U3/U6 promoters. Additionally, sgRNAs were expressed as polycistronic transcripts from Polymerase II (Pol II)-transcribed promoters, and mature sgRNAs are subsequently produced by cleavage through Csy4, self-cleaving ribozymes or the endogeneous tRNA processing system (Gao and Zhao, 2014; Nissim et al., 2014; Xie et al., 2015; Tang et al., 2016). Different Pol III promoters and Pol II-driven ribozyme, tRNA and Csy4 systems were recently systematically compared for genome editing in tomato protoplasts. The Csy4 and tRNA systems improved genome editing efficiencies approximately two-fold in comparison to Pol III-driven sgRNAs (Cermak et al., 2017). However, similar as for the nuclease itself, Pol II promoters providing strong and timely expression in the embryo will most likely be required to improve RGN efficiency by these approaches in Arabidopsis. Recently, also an optimized sgRNA scaffold developed for mammalian cells was reported to enhance editing efficiencies in rice, and in particular the abundance of biallelic and double mutants increased (Dang et al., 2015; Hu et al., 2018). We have now also implemented this improved sgRNA backbone to our system, but are awaiting experimental data to confirm improved efficiency. It should be noted that Cermak et al. (2017) did not observe improved efficiencies when employing a similar optimized sgRNA architecture (Chen et al., 2013). Additional to optimizing components or expression of the RGN system itself, e.g. co-expression of the exonuclease Trex2 was reported to improve genome editing efficiencies ~ two-fold in tomato and barley protoplasts (Cermak et al., 2017). Exonuclease co-expression concomitantly augmented average deletion size, which may facilitate initial mutation detection and simplify design of genetic markers. The described approaches provide ample opportunities to further boost genome editing efficiencies in the Arabidopsis systems towards the development of an optimal reverse genetic tool. Based on our findings that *RPS5a* and egg cell specific promoters confer highest genome editing efficiencies, we propose that one of the well-characterized vector systems incorporating these elements (e.g. Wang et al., 2015; Tsutsui and Higashiyama, 2017; or as described here) should be included for any further benchmarking of Cas9-based RGNs in Arabidopsis.

### Author contributions

JS and JO conceived the work, performed experiments and analyzed data. CK performed experiments. MB, LD’O and RB conceived Lhcb1-related experiments, and MB performed experiments. JS wrote the manuscript with contributions from all authors.

## Acknowledgements

Acknowledgements

We acknowledge Bianca Rosinsky for taking care of plant growth facilities and growing plants. Ulla Bonas is acknowledged for generous support. This work was funded by GRC grant STU 642-1/1 (Deutsche Forschungsgemeinschaft, DFG) and seed funding by the CRC 648 (DFG) to Johannes Stuttmann. Mauro Bressan and Luca Dall’Osto were supported by University of Verona (Program CooperInt2017 and HuntingLight Ricerca di Base 2015).

**Online Resource 1**: Architecture and functional verification of an adaptable, GUS-based nuclease activity reporter

(a) T-DNA region of the adaptable reporter plasmid, and cloning of user-defined target sequences. The “empty” plasmid contains a 35S-driven *GUS*, with a BsmBI-excisable ccdB cassette inserted between the initiating ATG and the *GUS* coding sequence. In a *BsmBI* Golden Gate reaction, the ccdB cassette is exchanged for a user-defined target sequence introduced as hybridzed oligonucleotides. A configuration in which the reporter detects a −1 nt repair event (or e.g. −4 or +2 nt events) is shown as example. The reporter may be adapted to detect different events by varying the length of the introduced target site. It should be noted that the introduced target site may not, after repair, contain an in-frame STOP codon.

(b) Functional verification of the GUS-based reporter. Two different target sites were introduced into the adaptable plasmid to obtain Reporters 1/2, reporters were co-expressed with respective nucleases in *N. benth.*, and GUS activity visualized by X-Gluc 3 dpi. Nuclease/reporter combination 2 consistently showed stronger GUS activity.

**Online Resource 2**: Sequence details on nuclease and reporter constructs used in this study (annotated GenBank files)

---

**Supplemental Figure S1:**
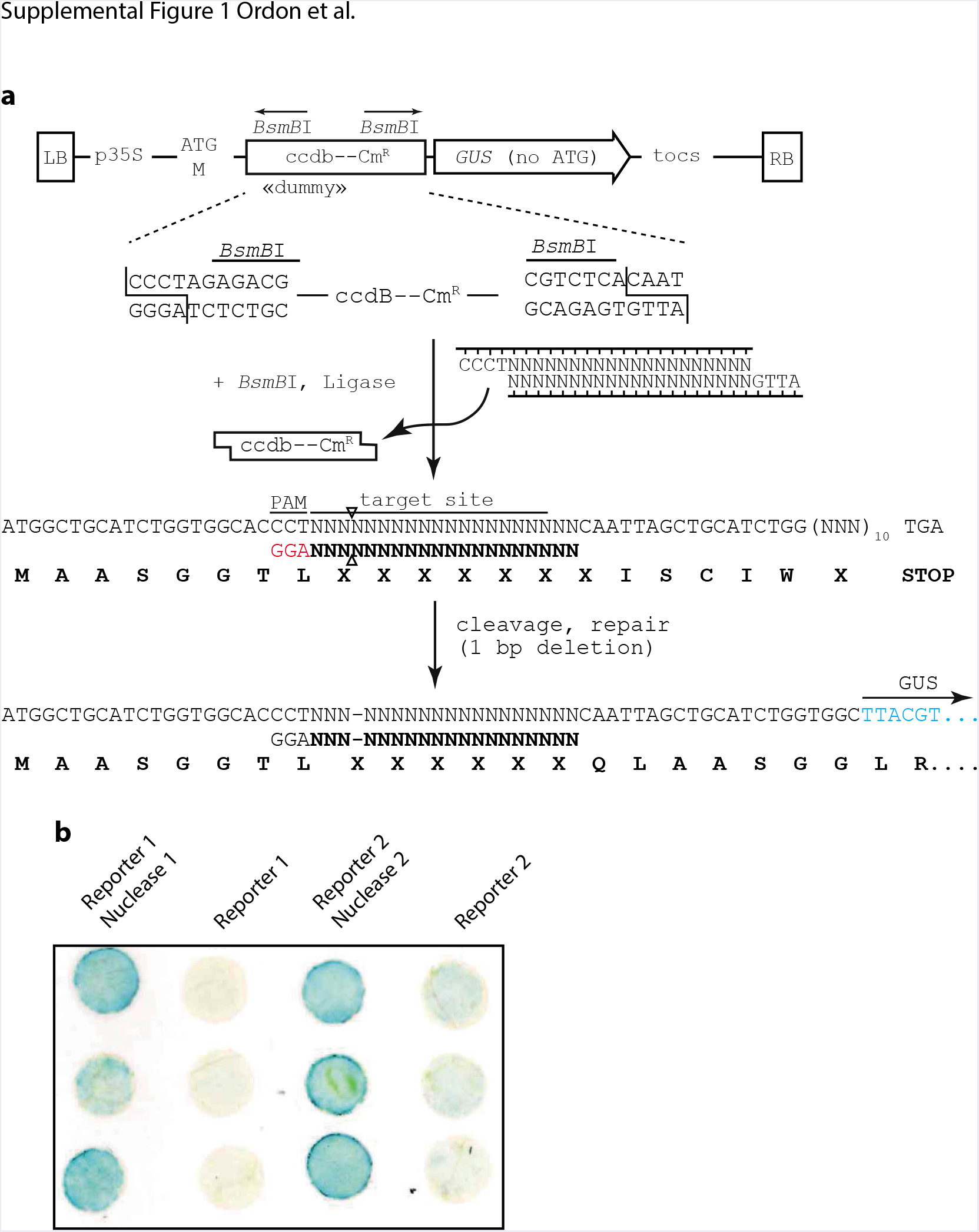
Architecture and functional verification of an adaptable, *GUS-based* nuclease activity reporter. (a) T-DNA region of the adaptable reporter plasmid, and cloning of user-defined target sequences. The “empty” plasmid contains a 35S-driven *GUS*, with a βsmβl-excisable ccdB cassette inserted between the initiating ATG and the *GUS* coding sequence. In a *BsmBI* Golden Gate reaction, the ccdB cassette is exchanged for a user-defined target sequence introduced as hybridzed oligonucleotides. A configuration in which the reporter detects a −1 nt repair event (or e.g. −4 or +2 nt events) is shown as example. The reporter may be adapted to detect different events by varying the length of the introduced target site. It should be noted that the introduced target site may not, after repair, contain an in-frame STOP codon. (b) Functional verification of the GUS-based reporter. Two different target sites were introduced into the adaptable plasmid to obtain Reporters 1/2, reporters were co-expressed with respective nucleases in *N. benth.*, and GUS activity visualized by X-Gluc 3 dpi. Nuclease/reporter combination 2 consistently showed stronger GUS activity.

